# Whisker map organization in somatosensory cortex of awake, behaving mice

**DOI:** 10.1101/587634

**Authors:** Han Chin Wang, Amy M. LeMessurier, Daniel E. Feldman

## Abstract

The whisker map in rodent somatosensory cortex is well characterized under anesthesia, but its organization during awake sensation, when cortical coding can differ strongly, is unknown. Using a novel behavioral task, we measured whisker receptive fields and maps in awake mice with 2-photon calcium imaging *in vivo*. During a whisker-attentive task, layer 2/3 pyramidal neurons were sharply tuned, with cells tuned to different whiskers intermixed in each column. This salt-and-pepper organization consisted of small clusters of similarly-tuned neurons superimposed on a mean subcolumnar map. Parvalbumin interneurons had broader tuning, and were more homogeneously tuned to the columnar whisker. During a sound-attentive task, whisker tuning of pyramidal cells was less heterogeneous in each column, and firing correlations increased. Thus, behavioral demands modulate fine-scale map structure, and decorrelate the whisker map during whisker-attentive behavior.

## INTRODUCTION

Layer (L) 2/3 provides the major cortical output of primary sensory cortex, but how sensory coding is organized in L2/3 is unclear. In rodents, salt-and-pepper intermixing of neurons tuned for different stimulus features is common, but the fine-scale organization, cell-type specificity, and behavioral state modulation of these intermixed maps are generally not understood. We studied these features in L2/3 of the whisker map in rodent somatosensory cortex (S1). L2/3 integrates feedforward, topographic sensory input from L4 with non-topographic input from secondary somatosensory thalamus, cross-columnar and other projections (Aronoff et al., 2010; Feldmeyer et al., 2013; Petersen and Crochet, 2013; Zhang and Bruno, 2019). In the classical, one-whisker-one-column model of S1, nearly all neurons in each cortical column are tuned to that column’s topographically matched whisker, termed the columnar whisker (CW) (Armstrong-James and Fox, 1987; Simons, 1978). But more recent cellular-resolution imaging, performed in anesthetized mice, revealed substantial salt-and-pepper intermixing of cells tuned for different whiskers in L2/3 of each column. This represents high local scatter overlaid on the gradual, average somatotopic gradient across L2/3 of S1 (Clancy et al., 2015; Margolis et al., 2012; Sato et al., 2007). Intermixing has been suggested to be locally random in S1 (Peron et al., 2015; Sato et al., 2007), unlike L2/3 of visual cortex (V1), which contains fine-scale clusters (5-40 µm) of similarly tuned neurons, which may represent columnar primitives (Kondo et al., 2016; Ringach et al., 2016; Rothschild et al., 2010).

Whether this organization also holds in S1 of awake animals, where cortical coding can differ strongly (Haider et al., 2013; Poulet and Petersen, 2008), is unknown. L2/3 is strongly influenced by brain and behavioral state through top-down input and neuromodulation (Petersen and Crochet, 2013). But whisker receptive fields have never been precisely measured in wakefulness, because active whisking prevents delivery of calibrated stimuli across whiskers in awake mice. Instead, prior studies have only roughly estimated whisker somatotopy from whisking onto objects, or by magnetically deflecting a single whisker (O’Connor et al., 2010; Peron et al., 2015; Pluta et al., 2017; Sachidhanandam et al., 2013). Thus whisker receptive fields, the organization of the whisker map, and their modulation by behavioral state are unknown in awake mice.

We developed a behavioral task that enables measurement of whisker receptive fields and the whisker map in L2/3 of S1 in awake, whisker-attentive mice. We characterized single whisker tuning and map organization of two major cell types—pyramidal (PYR) and parvalbumin (PV) neurons—using *in vivo* two-photon Ca^2+^ imaging. The awake map showed prominent salt-and-pepper organization that varied by cell type, contained local clusters of similarly-tuned neurons, and was influenced by sensory and attentional task features.

## RESULTS

### Whisker discrimination task for awake receptive field measurements

To characterize receptive fields in awake, whisker-attentive mice, we developed a whisker discrimination task that required mice to respond to specific whisker stimuli. Mice (n=10) were head-fixed, had 9 of their whiskers inserted in a 3 × 3 piezoelectric actuator array, and learned to lick to all-whisker stimuli (S+, simultaneous deflection of 9 whiskers), but not single-whisker stimuli (whisker S-), no-stimulus blank trials, or two auditory tones (tone S-) (Figure 1A). Mice learned this whisker-cued (WC) task, and false-alarm licking showed that they attended to whisker stimuli more than auditory distractors (Figure 1B-C). Mice moved their whiskers during <5% of whisker S− trials (Figure S1). GCaMP6s (Chen et al., 2013) was virally expressed in either L2/3 PYR or PV neurons using Emx1-Cre or PV-Cre mice, respectively, and stimulus-evoked activity was measured with two-photon Ca^2+^ imaging via a chronic cranial window (Figure 1D). Whisker receptive fields were measured only from whisker S− trials (Figure 1E-G), excluding lick (false alarm) trials, to avoid motion artefact and lick-related neural activity. Imaged neurons were localized relative to anatomical column boundaries via post hoc reconstruction of imaging fields and cytochrome oxidase staining of L4 barrels (Golshani et al., 2009).

**Figure. 1.**
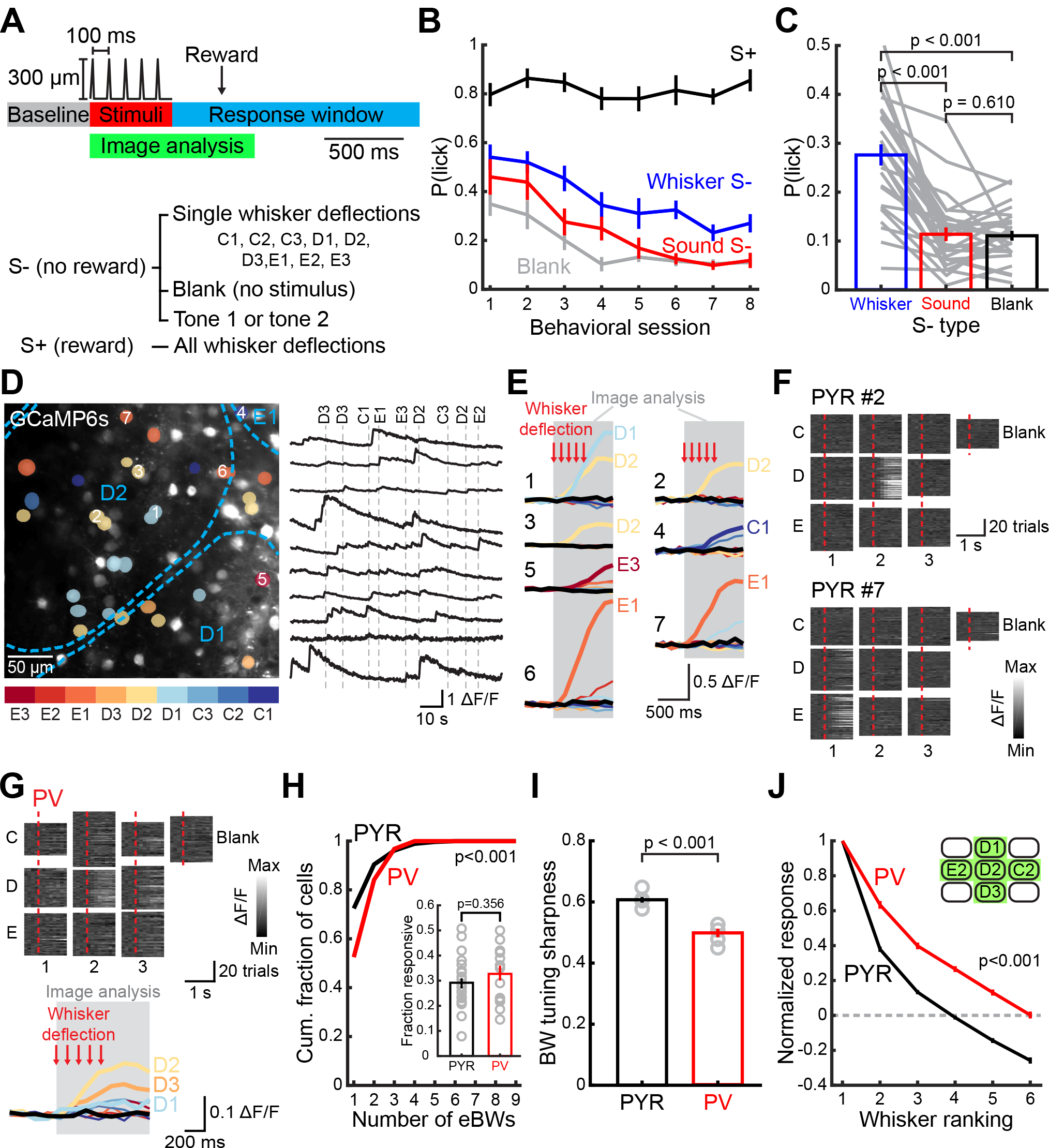
Whisker tuning of L2/3 PYR and PV neurons in awake, whisker-attentive mice. **(A)** Structure of a single trial (top) and stimuli-reward assignment for WC task (bottom). **(B)** Behavioral performance over 8 days before imaging. **(C)** Lick probability for different S-stimulus types during last 3 days in (B). Each gray lines represents one mouse in one session. Statistics: Friedman test. **(D)** Left: Example imaging field, showing barrel boundaries and cells color-coded for their BW. Right: Example ΔF/F traces from 10 cells. Dash: onset of stimuli. **(E)** Mean ΔF/F traces for 7 cells from the imaging field in (D). Color represents different whisker stimuli and blank (black); stimulus-labeled traces are significant responses. **(F)** ΔF/F traces for each whisker S− and blank trial, sorted by whisker stimulus, for 2 cells in (E). Dash, stimulus onset. Gray scale is normalized to the maximal response of that cell. **(G)** Example PV neuron with ΔF/F traces for each whisker S− and blank trial (top) and mean ΔF/F responses (bottom). Color represents different whisker stimuli and blank (black); stimulus-labeled traces are significant responses. **(H)** Receptive field size quantified as the number of eBWs. Statistics: χ^2^. (Inset) Fraction of whisker-responsive neurons in an imaging field. Each circle represents one imaging field. Statistics: rank-sum. **(I)** Mean tuning sharpness of responsive neurons. Gray circles show the mean for each animal. Statistics: rank-sum. **(J)** Mean rank-ordered tuning curve, after ranking stimuli within cells. Only cells located in green columns (inset) were included. Statistics: unbalanced two-way ANOVA. All error bars are SEM.

### L2/3 PYR neurons show heterogeneous, locally clustered tuning

Among PYR cells, 29% were whisker-responsive, as expected from sparse coding (Crochet et al., 2011; Estebanez et al., 2012; O’Connor et al., 2010; Peron et al., 2015). Whisker tuning was generally narrow. Of responsive cells, 72% had one best whisker (BW) that elicited a statistically stronger response than all other whiskers, and 45% responded only to the BW. The other 28% of responsive neurons had several statistically equivalent best whiskers (eBWs), indicating broadly tuned cells (Figure 1H). A tuning sharpness index (Figure 1I), which compared the response magnitude of BW to other whiskers, and mean rank-ordered tuning curves (Figure 1J) both reported generally sharp whisker tuning.

PYR neurons within each column were heterogeneously tuned (e.g., Figure 1D-E), similar to the salt-and-pepper organization under anesthesia (Clancy et al., 2015). The set of PYR neurons tuned to a given whisker (termed the tuning ensemble) spanned several columns, with only 36% of cells located within the whisker’s anatomical column (directly above the L4 barrel of that whisker, Figure 2A-B). Correspondingly, within each column only 45% of PYR neurons were tuned to the CW (Figure 2C). Each tuning ensemble was centered on its topographically appropriate S1 column, reflecting the average columnar organization of S1 (Figure 2A and D). Within each column, the neurons that were tuned to other nearby whiskers formed a rough subcolumnar map (Figure S2E). Thus, salt-and-pepper tuning heterogeneity is overlaid on the classical somatotopic whisker map, which is evident both across and within columns. The tuning preference for a reference whisker fell off steadily with PYR cell’s distance from the reference column center (Figure 2F). This occurred without a sharp boundary at the column edge, confirming a gradual mean tuning gradient within and across S1 columns (Sato et al., 2007).

**Figure 2.**
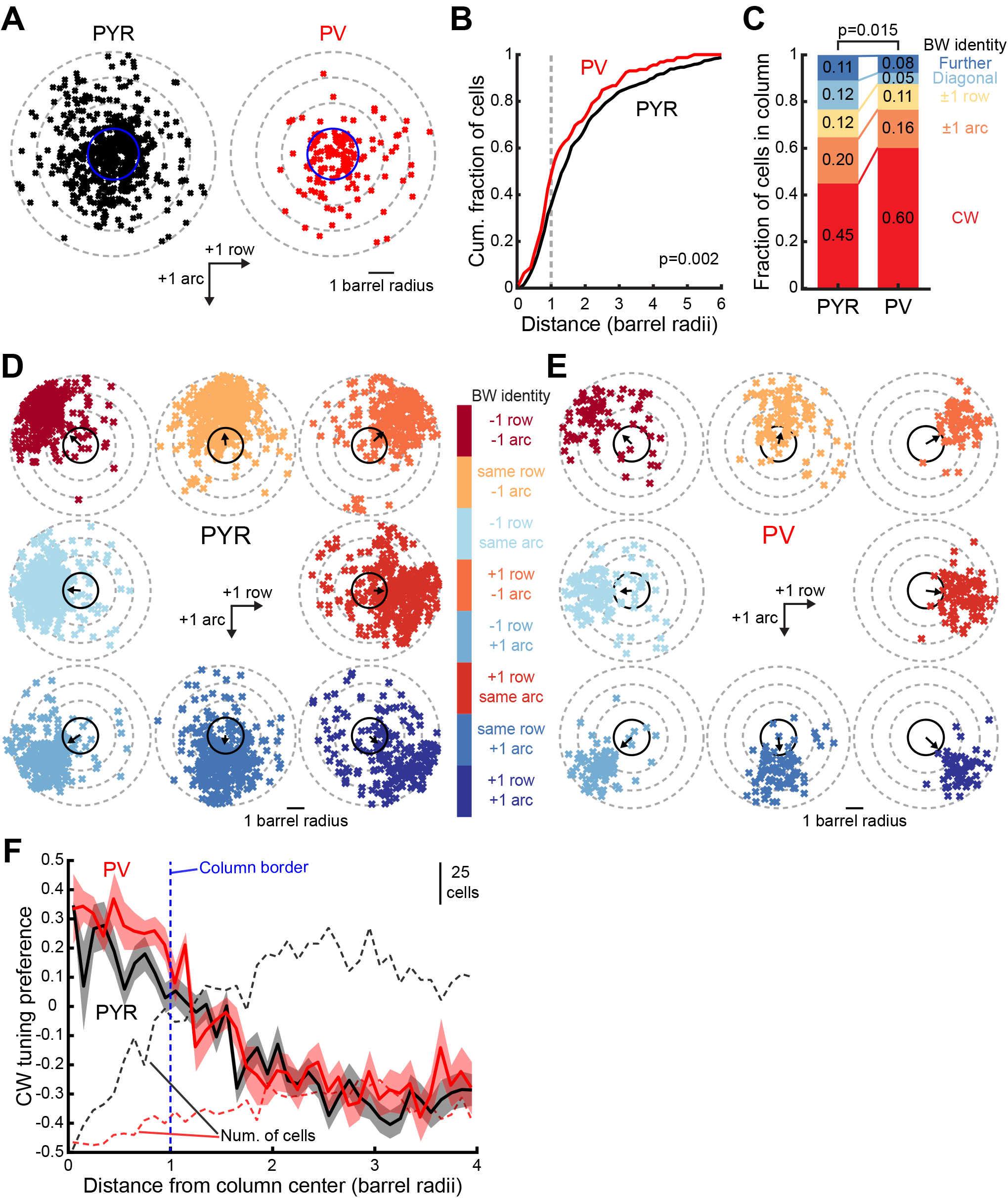
Whisker map topography for L2/3 PYR and PV neurons in whisker-cued mice. **(A)** Location of each responsive PYR and PV neuron relative to the anatomical column of its BW (blue circle). **(B)** Cumulative distribution of distance to BW column center for cells in (A). Statistics: KS. **(C)** Identity of BW for all responsive cells within a column. Statistics: χ^2^. **(D-E)** Location of PYR (D) and PV neurons (E) tuned to each of the 8 whiskers surrounding a reference whisker. The BWs of cells were color-coded, and the location of cells were plotted as their relative position to the reference whisker column (blank circle in the center). Arrows show circular mean of cell positions. **(F)** Tuning preference of responsive neurons for a reference whisker, as a function of distance from reference column center. PYR tuning shows a gradual gradient (Sato et al., 2007), while PV tuning shifts more sharply at column edge.

To determine if local clusters of similarly tuned neurons exist within the salt-and-pepper map, as in V1 (Kondo et al., 2016; Ringach et al., 2016), we examined pairwise signal correlations (tuning similarity) for co-columnar neuron pairs. Signal correlation fell off with distance between paired neurons (Figure 3A, red). Neurons < 30 µm apart had higher signal correlation than expected by chance, calculated by randomly shuffling neuron locations within each column (Figure 3A, gray line). To test whether this local tuning similarity simply reflects the average tuning gradient across S1, we estimated that mean tuning gradient in 2 dimensions by projecting all neurons in each anatomical column (across all imaging fields and mice) into one virtual column (Figure 3B). This virtual population will preserve the average sub-columnar tuning gradient, which is consistent across fields and mice by definition, but will abolish any local tuning similarity that does not occur in stereotyped columnar locations (see Methods). This analysis revealed the mean subcolumnar tuning gradient in within-arc and within-row dimensions, and showed that co-columnar PYR pairs < 30 µm apart have higher signal correlation than the mean subcolumnar tuning gradient (Figure 3C-F). Similar clustering was observed for 2 other measures of pairwise tuning similarity, including that nearby cells were 50% more likely to share the same BW than predicted by the average tuning gradient (Figures 3G-H, and S3). In contrast, there was no clustering for whisker response magnitude, which also varies across L2/3 PYR cells (Figure 3I). Thus, each S1 column contains a mosaic of PYR local tuning clusters, scattered around the mean subcolumnar map.

**Figure 3.**
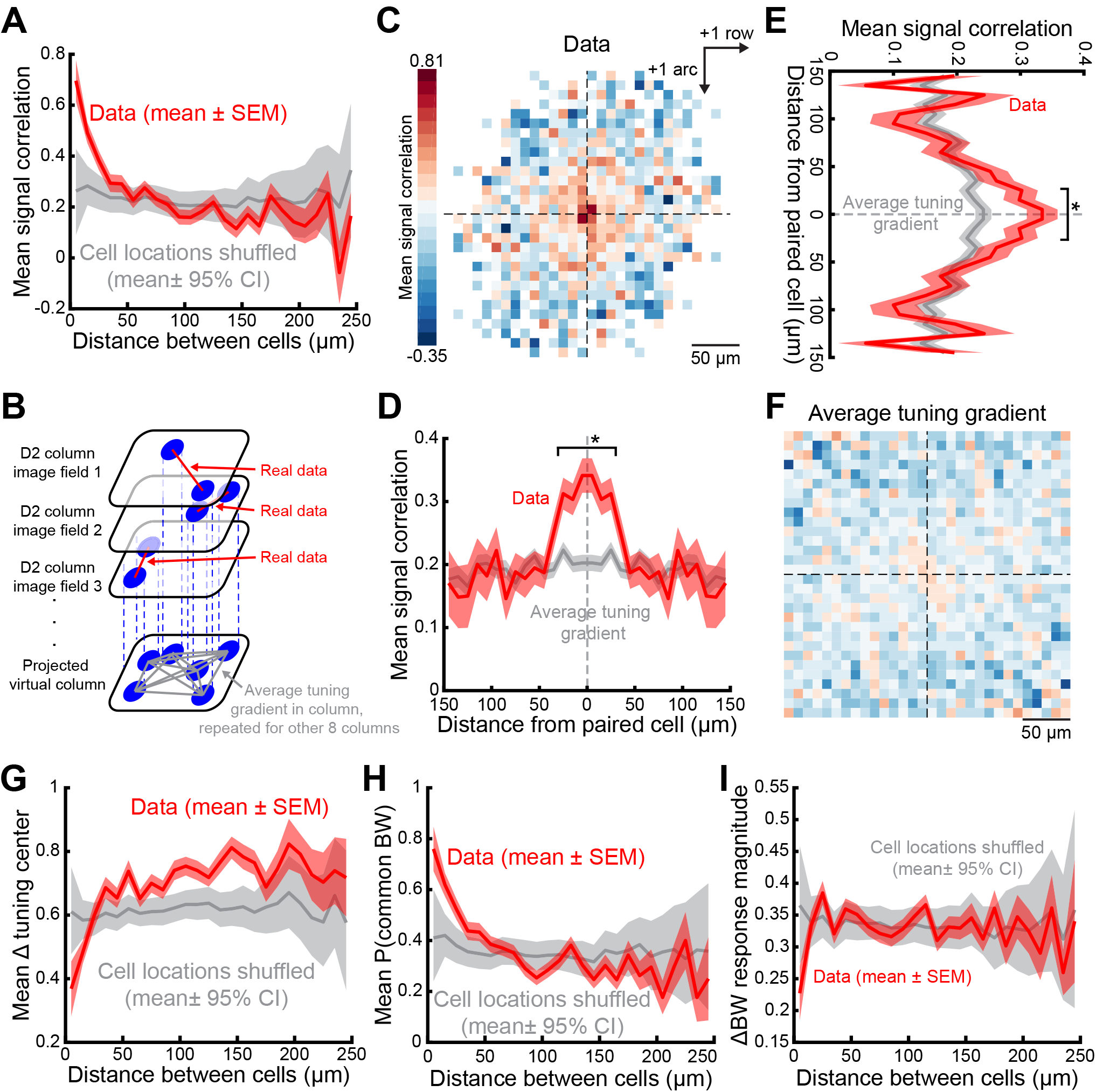
Similarly tuned neurons formed small clusters within whisker column. **(A)** Mean signal correlation for pairs of co-columnar PYR neurons, averaged within each distance bin, as a function of distance between neuron pairs. Red: observed data. Gray: spatially shuffled neurons. **(B)** Method to determine average sub-columnar tuning gradient in the absence of local clustering. **(C)** Mean signal correlation between co-columnar PYR pairs as a function of 2-D spatial position of cells. For each neuronal pair, one cell is positioned at the center, and the signal correlation between the pair is assigned to the spatial bin at the location of the other cell. Color shows the mean tuning difference within each 10 µm spatial bin with cell numbers in each bin ≥ 5 cells. **(D-E)**, Mean projection of data in (C) along row (D) or arc (E) dimension. *, significant difference for each bin relative to predicted column-average tuning gradient (gray). All shadings: SEM. **(F)** Predicted change of signal correlations in the same coordinate system as (C), with local clustering removed by projecting cells across imaging fields and mice. See Methods. **(G)** Mean difference in tuning center-of-mass for co-columnar PYR neuron pairs, as a function of distance between neuron pairs. Red: observed data. Gray: spatially shuffled neurons. Probability that a pair of co-columnar PYR neurons have the same BW, as a function of distance between neuron pairs. Red: observed data. Gray: spatially shuffled neurons. **(H)** Mean difference in normalized BW response magnitude for pairs of co-columnar PYR neurons, as a function of distance between neurons. Red: observed data. Gray: spatially shuffled neurons.

### Broader, more columnar tuning for PV neurons

PV interneurons powerfully shape information coding (Lee et al., 2012), and their activity varies with behavioral state (Pala and Petersen, 2018). In awake V1, PV neurons are often more broadly tuned than PYR cells (Hofer et al., 2011; Kerlin et al., 2010), but PV tuning in awake S1 is unknown. Imaging from L2/3 PV cells showed that PV cells had substantially broader whisker tuning than PYR cells. PV cells had more equivalently responsive whiskers (Figure 1H), and a broader rank-ordered whisker tuning curve (Figure 1I-J). At the population level, most PV neurons (60%) in a column were tuned to the CW (Figure 2A-C), yielding a more precise columnar map than for PYR cells (Figure 2E). Broad, CW-centered tuning by PV cells may reflect averaging of inputs from nearby, heterogeneously tuned PYR cells (Runyan et al., 2010). Unlike PYR cells, whose mean whisker tuning preference declined gradually with distance from each whisker column center, PV cells showed more homogeneous tuning within a column, which fell off more steeply at the column edge (Figure 2F). Thus PV cells exhibited a more pronounced columnar topography. Consistently, PV cells overlying L4 septa had slightly broader tuning than those overlying L4 barrels, by some measures, which was not observed for PYR neurons (Figure S2A-D). PV cells did not show local tuning clusters (Figure S3).

### Map differences between whisker-attentive and sound-attentive mice

L2/3 of S1 exhibits prominent experience-dependent plasticity (Feldman and Brecht, 2005; Margolis et al., 2014). To test if our behavioral training paradigm shaped the map organization described above, additional mice (n=9) were presented with the identical stimulus set, but were trained to lick to tone 1, but not to whisker stimuli (Figure 4A). This is termed the sound-cued (SC) task. False-alarm licking confirmed that mice attended to auditory stimuli instead of whisker stimuli (Figure 4B-C). PYR cell response amplitude and tuning sharpness were identical in WC and SC mice (Figure 4D-F). However, SC mice showed less salt-and-pepper tuning heterogeneity, evidenced by a spatially narrower tuning ensemble and an increased fraction of PYR cells tuned for the CW (Figures 4G-H). As a result, signal correlations between co-columnar neurons were substantially higher in SC mice (Figure 4I). In addition, noise correlations (trial-by-trial co-variability, Kohn et al., 2016) between pairs of co-columnar neurons were also higher in SC mice (Figure 4J). Elevated noise correlations were also found specifically for co-tuned neuron pairs in SC mice (Figure 4K), which is a condition that reduces sensory coding efficiency (Averbeck et al., 2006). These effects on signal and noise correlations were confirmed after sub-sampling to ensure identical spatial distributions of neurons in S1 (Figure S4A-G). Thus, maps in WC and SC are grossly similar, but local tuning and shared activity fluctuations are significantly decorrelated in the WC task. This may reflect training history, or acute attention to whisker stimuli (Harris and Thiele, 2011). No substantial differences in PV neuron properties were found between WC and SC groups (Figure S4H-N).

**Figure 4.**
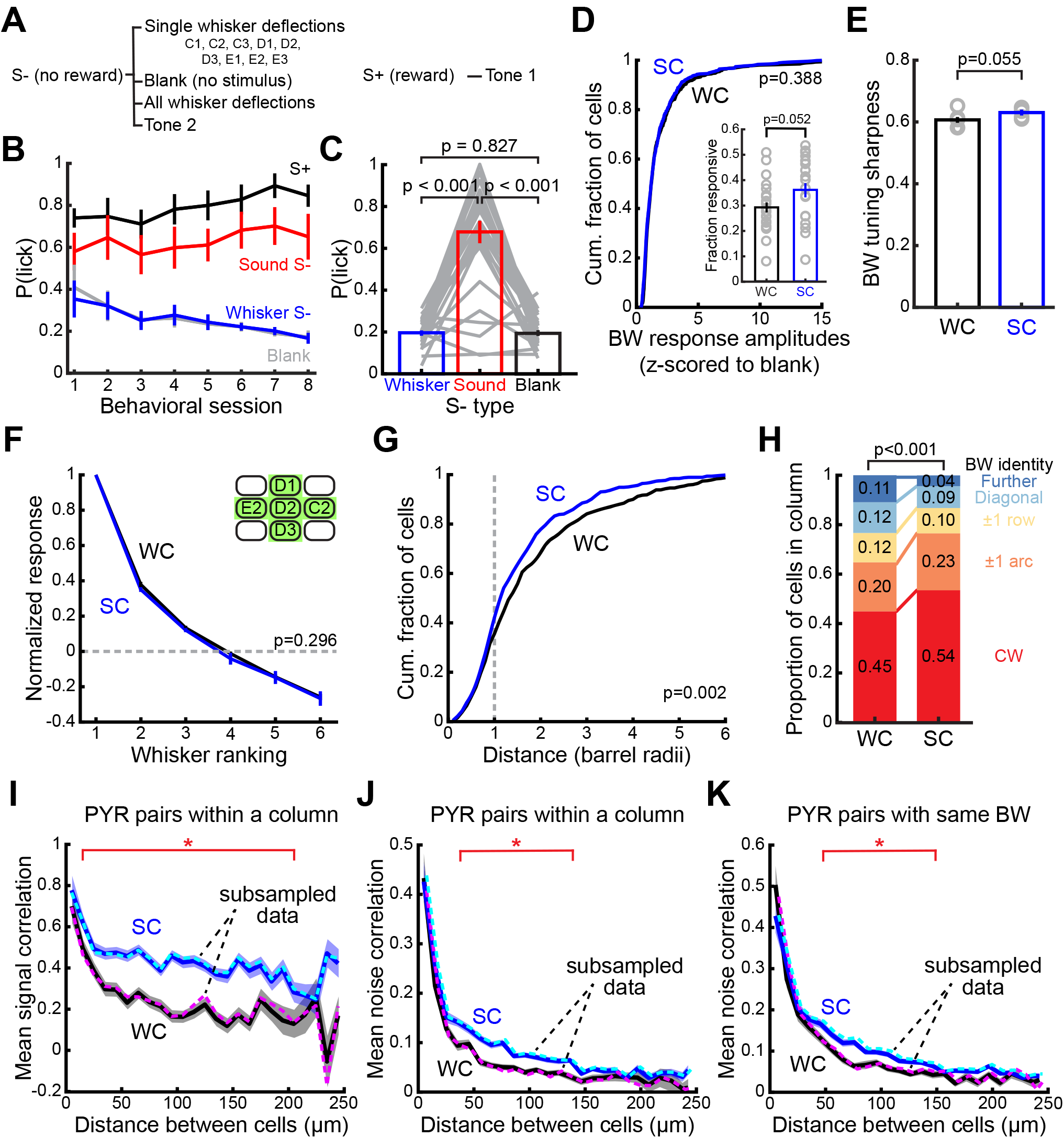
Whisker map differences between whisker-cued and sound-cued tasks. **(A)** Stimulus-reward assignment for SC task. **(B)** Behavioral performance over 8 days before imaging. **(C)** Lick probability for different S-stimulus types during last 3 days in (B). Each gray lines represents one mouse in one session. Statistics: Friedman test. **(D)** Normalized BW response magnitude for WC and SC PYR cells. Statistics: KS. (Inset) Fraction of whisker-responsive neurons in an imaging field. Each circle represents one imaging field. Statistics: rank-sum. **(E)** Mean tuning sharpness of responsive neurons. Gray circles show the mean for each animal. Statistics: rank-sum. **(F)** Mean rank-ordered tuning curve, after ranking stimuli within cells. Only cells located in green columns (inset) were included. Statistics: unbalanced two-way ANOVA. **(G)** Cumulative distribution of distance from responsive cells to their BW column center. Statistics: KS. **(H)** Identity of BW for all responsive cells within a column. Statistics: χ^2^. **(I)** Mean signal correlation and **(J)** noise correlation for pairs of co-columnar PYR neurons, averaged within each distance bin, as a function of distance between neuron pairs. Dash: mean correlations after subsampling neurons to ensure identical columnar distributions in WC and SC. **(K)** Same as (J), but for PYR pairs with the same BW. *, significant difference between SC and WC, permutation test within each bin. All error bars or shading are SEM.

## DISCUSSION

Thus, awake S1 contains a globally ordered but locally heterogeneous whisker map among L2/3 PYR cells, resembling awake V1 (Bonin et al., 2011; Ohki et al., 2005) and A1 (Rothschild and Mizrahi, 2015). Within each column, individual PYR cells were sharply tuned for one or a few whiskers, but cells tuned for the CW and nearby whiskers were locally intermixed. This was quantitatively similar to L2/3 organization in anesthetized mice (Clancy et al., 2015). Within this salt-and-pepper organization, local clusters of similarly tuned neurons exist. This noisy, clustered micro-organization is superimposed on the average somatotopic tuning gradient that exists within and across S1 columns (Armstrong-James and Fox, 1987; Sato et al., 2007). L2/3 also contains a noisy gradient of angular tuning (Andermann and Moore, 2006; Bruno et al., 2003; Kremer et al., 2011; Peron et al., 2015), roughness tuning (Garion et al., 2014), and domains that vary in single-versus multi-whisker stimulus preference (Estebanez et al., 2016; Estebanez et al., 2012). Thus, clustered salt-and-pepper organization may be a byproduct of compressing multiple high-dimensional feature representations into limited space in each column.

Prior studies of S1 in anesthetized mice did not observe spatial clustering of functionally similar PYR neurons in excess of average columnar tuning gradients (Bruno et al., 2003;, but see Garion et al., 2014; Kerr et al., 2007; Martini et al., 2017; Peron et al., 2015; Sato et al., 2007). One study in awake mice also found functional clustering to be absent, and concluded that response properties were randomly intermixed in L2/3 (Peron et al., 2015). This study did not examine somatotopic tuning. In contrast, we observed pronounced local clustering on the <30 µm scale within each column, in which nearby neurons were more likely to share whisker tuning than predicted by chance, or than expected from the subcolumnar tuning gradient. Thus S1 appears similar to rodent V1, which contains microcolumn-like structures of co-tuned neurons on the 5-40 µm scale (Kondo et al., 2016; Ringach et al., 2016). Such clusters have not yet been found in auditory cortex (Bandyopadhyay et al., 2010; Issa et al., 2014; Panniello et al., 2018). Clustering may enable more efficient sensory computation via local synaptic connections, and may develop from shared connectivity of developmentally related neurons (Li et al., 2012), or activity-dependent selection of common inputs due to shared local connectivity.

The basic organization of the L2/3 whisker map was similar in WC and SC mice, and thus was not a product of specific training on the whisker-attentive task. Mice performing the WC task showed slightly more salt-and-pepper tuning heterogeneity in each column, and lower tuning similarity and noise correlations between co-columnar neurons. Thus, sensory tuning and neural activity were decorrelated within the neural population of each column in the WC task. This decorrelation is expected to provide better whisker discrimination and detection performance by columnar ensembles in the whisker-attentive task (Averbeck et al., 2006; Kohn et al., 2016). The difference between the WC-versus SC-trained mice could reflect either training history or acute effects of attending to whiskers or sound, which were not distinguished in this study. This demonstrates that activity and fine structure of the clustered, salt-and-pepper map in L2/3 are mutable to support task demands, as in V1 of mice and primates (Bondy et al., 2018; Goltstein et al., 2018; Ramalingam et al., 2013).

These results demonstrate a marked transformation of whisker sensory representation in L2/3 compared to L4, which has highly accurate columnar topography in awake mice (Hires et al., 2015). Salt-and-pepper intermixing of neurons tuned to different whiskers may reflect divergence of L4-L2/3 projections, cross-columnar spread of information in L2/3, feedback from secondary somatosensory cortex, and/or integration of multi-whisker thalamic input (Aronoff et al., 2010; Bender et al., 2003; Feldmeyer et al., 2013; Kwon et al., 2016; Yang et al., 2016; Zhang and Bruno, 2019). Real-world tactile input is inherently multi-whisker, so local intermixing of neurons tuned to different whiskers could be an intermediate step in synthesis of precise multi-whisker feature tuning (Ramirez et al., 2014). While serial transformations in sensory population codes typically emerge across distinct cortical areas in large mammals, the small size of the rodent brain may compress multiple representations within each cortical area. How this spatially intermixed code is read out by downstream projection targets of L2/3 PYR cells remains unclear.

## AUTHOR CONTRIBUTIONS

H.C.W. and D.E.F. designed the study. H.C.W. performed the experiments and analyzed the data. A.M.L. provided analysis software. H.C.W. and D.E.F. wrote the manuscript.

## Supplementary materials and methods

### Animals

Animal procedures were approved by the UC Berkeley Animal Care and Use Committee, and followed NIH guidelines. Mice were obtained from The Jackson Laboratory and bred in C57BL/6J background. 11 Emx1-IRES-Cre mice (JAX 005628) were used, 6 for whisker-cued experiments and 5 for sound-cued experiments. 8 PV-IRES-Cre mice (JAX 017320) were used, 4 for whisker-cued experiments and 4 for sound-cued experiments. Mice were of either sex.

### Surgery and viral injection

At 2.5–3 months of age, mice were anesthetized with isoflurane (1-1.5% in O_2_), and a stainless steel head holder containing a 6 mm diameter aperture was affixed to the skull over S1 using cyanoacrylate glue and dental cement (C&B Metabond, Parkell). The location of D1, D2 and D3 whisker columns was mapped using transcranial intrinsic signal optical imaging (Drew and Feldman, 2009; Grinvald et al., 1986). These columns correspond to the central row of whiskers stimulated by our 3-by-3 piezo actuator array. A 3 mm diameter craniotomy was made centered on the D2 column. AAV viral vectors encoding Ca^2+^ indicator GCaMP6s (AAV2/1.Syn.Flex.GCaMP6s.WPRE.SV40, UPenn Vector Core AV-1-PV2821) were injected at 3-4 locations surrounding the D2 column at two depths (250 µm and 350 µm below pial surface). Injection was via nanoliter injector (Nanoliter 2000, World Precision Instruments). The craniotomy was covered with a chronic cranial window consisting of a 3 mm diameter glass coverslip (#1 thickness, CS-3R, Warner Instrument) sealed by dental cement. Before surgery, mice received dexamethasone (2 mg/kg), meloxicam (10 mg/kg) and enrofloxican (5 mg/kg) subcutaneously to reduce inflammation, provide prophylactic analgesia, and prevent infection. Following surgery mice were given buprenorphine (0.1 mg/kg) subcutaneously for analgesia.

### Behavioral task

One behavioral training session was performed each day. At the start of each session, the mouse was transiently anesthetized with isoflurane and head-fixed under the 2-photon microscope. 9 whiskers (rows C-E, arcs 1-3) were inserted into a 3 × 3 array of calibrated piezoelectric actuators (PSI-5H4E, PiezoSystems) controlled by a piezo amplifier (E503, Physik Instrumente). Whiskers were not trimmed, and were threaded through a tube mounted on each piezo with the tip of tube 5 mm from the whisker pad, and held in place by soft glue. A plastic shield prevented piezos from contacting other whiskers. A drink port with capacitive lick sensor was placed in front of the snout. Paw guards prevented paw contact with the whiskers, piezos, or drink port. Mice woke up from anesthesia and performed the behavioral task.

The task used a trial-based structure. Each trial lasted 2.5 s, consisting of a 0.5 s baseline period, 0.5 s stimulus period, and 1.5 s response window. During the stimulus period, 1 out of 12 possible stimuli were presented (9 single whisker deflections, all-whisker deflection, and 2 tones) or no stimulus (blank). Whisker stimuli consisted of 300 µm amplitude (equivalent to 6⁰ angular deflection) rostro-caudal, ramp-and-return deflections with 2 ms rise/fall time and 10 ms duration. A train of 5 deflections at 100 ms interval was used to increase the probability of multi-spike responses which would more reliably evoke GCaMP signals. Single-whisker deflections were delivered to one randomly chosen whisker. All-whisker deflections were simultaneous across all 9 whiskers. Tone stimuli were a single tone pip (2 or 8 kHz, 200 ms duration), delivered from a speaker 10 cm from the mouse. All stimuli and blank trials were presented in a pseudo-random order. The inter-trial interval was a 3 ± 1 s, after which mice needed to suppress licking below a threshold to initiate the next trial. Thus, consecutive stimuli were separated by at least 5.5 ± 1 s, which minimizes adaptation and short-term plasticity.

On S+ trials, water reward (2-4 μl) was delivered 300 ms into the reward window (i.e., 1300 ms after trial start, Figure 1A). Water was not delivered during S− trials. Licking above a threshold rate during the response window was scored as a lick response. No punishment was given for false alarm or miss trials. Licking was not required for water delivery, but during training, mice learned that S+ stimuli predicted reward, and learned to lick prior to reward delivery time (see ‘Training stages’).

Training was performed in total visual darkness (using 850 nm IR illumination for behavioral monitoring camera), and background white noise masked sounds from the piezo actuators. Tasks were controlled by an Arduino Mega 2560 microcontroller (Arduino) under user control and monitoring from custom routines in Igor Pro (WaveMetrics).

### Training stages

Initial training began 1-2 weeks after cranial window implantation. Mice were water-restricted to 85% of ad lib body weight, with free access to food. Daily water intake (0.7-1.0 mL) was calibrated individually for each mouse, and weight and health were monitored daily. Training proceeded in stages. In Stage 1, mice were acclimated to head-fixation and the presence of the water port. Paw guards prevented forelimb reaching to the whiskers or water port. In Stage 2, mice learned to lick for water rewards (2-4 μl) cued by simultaneous blue light flashes from an LED mounted on the drink port. In Stage 3, S+/S− training was begun using all-whisker deflection, one tone stimulus, and blank trials. For WC mice, the all-whisker stimulus was the S+. For SC mice, the tone was the S+. The other stimulus and blank were not rewarded. The blue light cue signaled water reward, which was presented 300 ms into the response window on S+ trials. Learning progress was evaluated by the fraction of false alarms and by a shift in lick timing from late in the response window (following the blue light cue) to early in the response window (prior to the blue light cue, indicating that the mouse was responding to the S+ stimulus itself, **Figure S1C-D**). This training stage continued until false alarm rate fell below 50%, and >50% of licks occurred prior to the blue light cue. In Stage 4, the final full behavioral task was implemented by introducing the other 10 S− stimuli and removing the blue light cue.

Imaging sessions began when mice reached stable performance in Stage 4, and the total number of trials per session exceeded 900 trials with about 20% S+ trials. All whiskers remained intact throughout the experiment.

### Two photon imaging

2-photon imaging was performed during behavior 4-6 weeks after viral injection. Imaging was performed with a Moveable Objective Microscope (MOM, Sutter) and Chameleon Ultra Ti:Sapphire mode-locked laser (Coherent). GCaMP6s was excited at 920 µm. Fast X-Y laser scanning was achieved by combining one resonant scanner (RESSCAN-MOM, Sutter) and one galvo scanner (Cambridge Technology). Green fluorescent emission was collected through a water-dipping objective (16X, 0.8 NA, N16XLWD-PF, Nikon), bandpass-filtered with a dichroic mirror (HQ 575/50, Chroma), and detected with GaAsP photomultiplier tubes (H10770PA-40, Hamamatsu). Laser power at the front of objective was 30-75 mW. Serial single plane images (512 × 512 pixels, corresponding to 305 µm × 305 µm, 175-250 µm below dura) were acquired at 7.5 Hz (30 Hz acquisition and averaging every 4 frames) using ScanImage5 software (Pologruto et al., 2003, Vidrio Technologies). For PV neurons, a larger imaging field was used (512 × 512 pixels, 406 × 406 µm), to sample more PV neurons. Imaging was performed daily. Each session comprised 900-1000 trials. 2-4 imaging fields were sampled in each mouse. In total, we obtained 24 imaging fields from whisker-cued Emx1-IRES-Cre mice, 20 from sound-cued Emx1-IRES-Cre mice, 14 from whisker-cued PV-IRES-Cre mice, and 15 from sound-cued PV-IRES-Cre mice.

### Histology and imaging field localization

Imaged cells were localized post-hoc relative to column boundaries defined by cytochrome oxidase staining of L4 barrels. To do this, an image z-stack was collected from the L2/3 imaging plane to the surface blood vessels at the end of each imaging session. After experiments were complete, the mouse was sacrificed and the brain was removed and fixed in 4% paraformaldehyde. The cortex was flattened and sectioned at 50 µm parallel to the cortical surface on a freezing microtome. Sections were processed for cytochrome oxidase staining, which reveals barrels in L4 and surface blood vessels. Imaging fields were localized relative to barrel column boundaries using surface blood vessels for alignment in ImageJ (NIH). Barrel centroids and the X-Y coordinate of each ROI relative to whisker column boundaries were calculated using custom Matlab (MathWorks) routines. A cell was considered to be within a column if more than half of its ROI pixels were located inside the L4 cytochrome oxidase boundary for that column. Other cells were classified as located overlying a septum. Barrel and septal cells were combined for analysis, except in figure S2A-D. In figures 1J and 4F, columns were defined by the mid-points of septa, so both barrel and septal cell were included.

### Calcium imaging analysis

Data analysis was conducted using custom routines in Matlab unless stated otherwise.

#### Imaging processing and ROI selection

Raw imaging data was corrected for slow X-Y motion in Matlab using NoRMCorre (Pnevmatikakis and Giovannucci, 2017). Substantial movement in Z axis was not observed, and not corrected. Neuronal regions-of-interests (ROIs) were defined manually from averaged frames as ellipsoid regions over neuronal somata. The fluorescence signal from each ROI (F_ROI_) was calculated as the average of its constituent pixels. For each ROI, neuropil contamination was subtracted by defining a neuropil ROI consisting of an annulus of 2-12 pixels outside the soma ROI border. Neuropil pixels that were spatially contained within other soma ROIs, or whose activity was correlated to any soma ROI (Pearson correlation coefficient > 0.2, Peron et al., 2015), were excluded from the neuropil ROI. Fluorescence of each soma ROI was then calculated as F_corrected_ = F_ROI_ − F_neuropil_, where F_neuropil_ is the average fluorescence in the neuropil ROI. F_corrected_ was constrained to be non-negative on each frame. Soma ROIs were excluded from analysis if F_ROI_ < F_neuropil_ in more than 0.5% of frames in any movie, which often represents bright or active neurites in the neuropil ROI. ΔF/F was calculated as (F_corrected_ − F_0_)/ F_0_, where baseline fluorescence F_0_ was defined as the 25^th^ percentile of F_corrected_ within a 60-sec sliding window.

For analysis, ROIs were required to be located ≤ 1.25 barrel radii from the centroid of one of the 9 stimulated whisker columns. Total numbers of neurons imaged in each cell type/task type were: whisker-cued PYR neuron: 2270; whisker-cued PV neuron: 499; sound-cued PYR neuron: 1839; sound-cued PV neuron: 424.

#### Whisker-evoked responses and receptive fields

Stimulus-evoked ΔF/F was measured on each trial as the mean ΔF/F for 1000 ms after stimulus onset minus mean ΔF/F for 500 ms pre-stimulus baseline. The mean whisker response of each ROI was then quantified across all non-lick S− trials. To identify significant whisker responses, we used a permutation test for difference in mean response relative to blank trials. In each iteration of this permutation test, single-trial ΔF/F data were shuffled between whisker S− and blank trials, and the difference in mean response between these shuffled trial sets was calculated. This was repeated 10,000 times to generate a null distribution. A measured whisker response was considered significant if it was > 95^th^ percentile of this null distribution. P-values were corrected for multiple comparisons across all S− stimuli with false discovery rate 0.05 (Benjamini-Hochberg procedure; Benjamini and Hochberg, 1995). A cell was considered whisker-responsive if at least one whisker induced a significant response.

Previous *in vivo* imaging studies (Peron et al., 2015) have shown that whisker stimulation can drive a reduction in GCaMP fluorescence below baseline (a negative ΔF/F response) in some cells, likely reflecting surround inhibition (Derdikman et al., 2003). In our data set, 11.6% of whisker-responsive PYR neurons (i.e., cells with a significant positive ΔF/F response to at least one whisker) and 9.5% of whisker-responsive PV neurons had a significant whisker-evoked negative ΔF/F to a non-BW whisker. Negative ΔF/F responses were small and slow compared to the more rapid, clearly time-locked excitatory responses. Negative ΔF/F responses were included when analyzing signal/noise correlations and rank-ordered tuning curve shape, but were excluded for calculation of tuning sharpness (see ‘Tuning of individual neurons’ and ‘Signal and noise correlation analysis’). In addition, 11.1% of PYR neurons and 2% of PV neurons exhibited only negative, not positive, significant ΔF/F responses (i.e. only inhibition, no excitation). These cells were considered non-whisker responsive and not analyzed further, because we could not rule out that they were excited by more distant, non-sampled whiskers.

#### Tuning of individual neurons

The best whisker (BW) was defined as the whisker that evoked the largest mean ΔF/F response and was significantly greater than blank trials. Equivalent best whiskers (eBWs) were defined as whiskers whose response amplitudes were not statistically different from the BW by permutation test, and were significantly greater than blank trials. BW tuning sharpness was defined as:

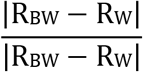

where R_BW_ = mean ΔF/F to BW, and R_W_ = averaged mean ΔF/F for all other whiskers (excluding any whiskers that evoked a negative ΔF/F response). Columnar whisker (CW) preference (Figure 2F) was calculated similarly as (R_CW_ - R_W_)/(R_CW_ + R_W_), where R_CW_ = ΔF/F to the CW. Rank-ordered tuning curves were calculated by ranking each stimulus from strongest to weakest within each cell, normalizing to the strongest response for that cell, and then averaging ranked tuning curves across cells. This quantifies tuning sharpness around each cell’s BW, independent of somatotopic organization of the receptive field. For rank-ordered tuning curves, only cells located at the center or one of the center-edges of the piezo array were included (Figures 1J and 4F, inset). This ensures that the BW plus 5 or 8 immediate adjacent whisker responses were sampled.

#### Normalized anatomical reference frame for spatial analysis across imaging fields

To combine imaging results across different imaging fields, ROI coordinates were transformed into a common reference frame in polar coordinates. This was done by first drawing a vector from the centroid of a reference column to the ROI. The normalized distance from ROI to column center was calculated as (measured distance) / (distance from column center to column edge along this vector). This gives units of barrel column radii. To determine the angular position for each ROI, vectors were drawn connecting the centroid of each surrounding column to the centroid of the reference column. These vectors were considered to be 45 degrees apart, i.e. evenly distributed around the reference column. ROI angle was determined relative to these vectors. This coordinate frame preserves ROI location relative to adjacent row- and arc-columns.

#### Signal and noise correlation analysis

To quantify receptive field similarity between pairs of simultaneously imaged neurons, we calculated signal correlation as the Pearson’s correlation coefficient between the mean responses to each single whisker stimulus. Noise correlations quantify the trial-by-trial co-variability between neuron pairs. To calculate this, for each single-whisker stimulus and blanks, a vector of single-trial responses (minus the mean response) was calculated, and the Pearson’s correlation coefficient was calculated between these vectors. Noise correlation for a neuron pair was the average correlation coefficient across 9 whiskers and blanks. Signal and noise correlations were averaged across cell pairs in 10 µm spatial distance bins. Significant differences in mean correlation values between WC and SC groups in each bin were determined by permutation test, relative to a null distribution in which correlation values were shuffled between WC and SC groups. P-values were corrected for multiple comparisons across all bins with false discovery rate of 0.05.

#### Assessment of spatial clustering of similarly tuned neurons

To test whether neurons are clustered by tuning properties, we calculated three measures of tuning similarity for pairs of simultaneously imaged neurons—signal correlation (Figure 3A-F), difference between center-of-mass of whisker tuning (Figures 3G and S3A-D), and probability of sharing the same BW (Figures 3H and S3E-H). To calculate each neuron’s center-of-mass of whisker tuning, whiskers were assigned to Cartesian coordinates in a 3 × 3 grid, and center-of-mass was calculated from the response amplitude to each whisker. These measures assess slightly different aspects of tuning: Signal correlation is influenced by all whiskers equally; tuning center-of-mass primarily reflects the strongest whiskers within the receptive field, and BW identity assesses only the strongest single whisker. We also measured similarity in general sensory responsiveness, by quantifying the relative difference in BW response strength between neurons as:

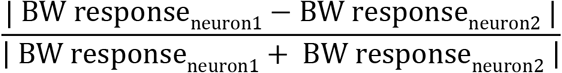

We calculated each measure for all simultaneously imaged, co-columnar pairs of PYR neurons, as a function of distance between cells in 10 µm spatial bins. Averaging within each bin revealed the mean tuning similarity as a function of distance between cells (red lines in Figure 3A and G-J). To test for significant spatial clustering relative to a null model of completely random intermixing of tuning properties, we compared these results to a shuffled dataset in which the locations of co-columnar neurons were randomly shuffled within each column, repeated 10,000 times to generate a null distribution (gray lines in Figure 3A and G-J). To assess clustering in 2 dimensions, we calculated the spatial position of each neuron (N1) relative to the other neuron in the pair (N0), in a pair-wise spatial reference frame that represents anatomical direction from N0 to N1 (relative to row and arc dimensions, as in “Normalized Anatomical Reference Frame” above) and absolute distance between the neurons. Tuning similarity was assigned to N1’s coordinate, and binned across pairs using 10 µm 2-D spatial bins. Mean tuning similarity was analyzed and plotted for each bin with > 5 cells (Figures 3C and S3A and E). Spatial profiles of tuning similarity in the arc or row dimension were obtained by projecting the 2D data along that dimension (e.g., Figure 3D-E). PV neurons were analyzed separately using the same method.

To test whether tuning similarity of local clusters was greater than expected from the mean tuning gradient across column (Figure 2F), we used a different comparison procedure. To estimate the average tuning gradient between neuron pairs, we calculated the 2-D spatial position of each ROI in a column-centered reference frame relative to the barrel centroid, including absolute distance from the centroid, as mentioned above. We then projected all neurons in a given anatomical column (e.g., the D2 column) across all imaging fields and animals into one virtual column (Figure 3B). This virtual population will contain the average tuning gradient. We sampled virtual PYR pairs from this combined data set, and calculated tuning similarity for these virtual co-columnar pairs, tracking results within the pair-wise spatial reference frame. This process was repeated for each imaged column (D1, E1, etc.). This reveals the mean topographic tuning gradient, because this gradient is consistent across fields and animals, but abolished local spatial clustering, because these are non-stereotyped relative to columnar location after we scrambled the cell pairs (Figure 3F). Significant differences between measured tuning similarity for real co-columnar pairs and this virtual combined data set was determined by permutation test within each distance bin, corrected for multiple comparisons across all bins with false discovery rate of 0.05. This analysis revealed significant local clusters of similarly-tuned neurons, greater than expected from the mean tuning gradient, over a spatial scale of ∼30 µm (Figure 3D-E). Clustering was also apparent from the other tuning similarity measures (Figures 3G-H, and S3A-H), but not for BW responsiveness (Figure 3I) or PV neurons (Figure S3I-K).

#### Active whisker movement during behavior

To monitor active whisker movement during behavior, we imaged movement of whiskers contralateral to the piezo actuator array. The compact design of the piezoelectric array prevented imaging whiskers within the actuators. Whiskers were imaged at 15 Hz throughout the 2-photon imaging period. 850 nm light from an infrared LED array provided background illumination. Whiskers were identified as black silhouettes relative to this IR background illumination. Movement was quantified for two C-row whiskers (either C1-C2 or C2-C3) in each mouse. Whisker positions were tracked by the local minimum pixel brightness within a curved ROI drawn along the trajectory of whisker movement, ∼2 cm from whisker pad. The instantaneous velocity of each whisker was calculated as the first derivative of whisker position. K-means clustering of velocity values was used to identify a velocity threshold that separated active movement from non-movement within each movie. For each whisker, we calculated the percentage of non-lick whisker S− stimulus trials that contained whisker movement. This was averaged across the 2 whiskers to give a single value per mouse (n=5).

#### Spatial subsampling of ROIs

Results reported in the main figures are for all ROIs that were whisker responsive and located in or near the stimulated whisker columns, as defined above. To validate functional differences between WC and SC mice, we performed additional analysis that corrected for small differences in spatial distribution of imaged neurons between WC and SC mice, by subsampling data to generate spatially identical sampling in WC and SC populations. For subsampling, each ROI was assigned to its closest whisker column. In one iteration of subsampling, cells were randomly chosen from WC and SC mice so that the numbers of cells assigned to each whisker column were the same in WC and SC mice. Data analysis was done for these subsampled cells. We performed 1000 iterations of this subsampling. The resulting mean and 95% confidence intervals are reported in figure S4.

### Statistics

Statistical analysis methods are described in Figure Legends and Methods (above). We did not pre-determine sample size, and all tests were two-tailed except for permutation test. Assignment to WC and SC groups was not randomized, but instead were run as two sequential cohorts of mice. During imaging analysis, identification of ROIs was done blind to stimulus-evoked response properties. Analyses were conducted using single neurons as the unit N, with the following exceptions: In figure 1C and 4C, behavior is quantified by behavioral session (gray lines each show one mouse on one session). In figure 1H inset and figure 4D inset, we analyzed the proportion of responsive neurons per imaging field, with each gray circle representing one imaging field. In figures 1I, 4E, S2C, and S4I, analyses were conducted on single neurons, but gray circles show the mean for each animal, to demonstrate consistency across mice.

Abbreviations for statistical methods are: KS, Kolmogorov-Smirnoff test. χ^2^, chi-squared test. Rank-sum, Wilcoxon rank-sum test. Data are presented as mean ± standard error of mean (SEM), except mentioned otherwise.

## Supplementary figure legends

**Supplementary figure 1.**
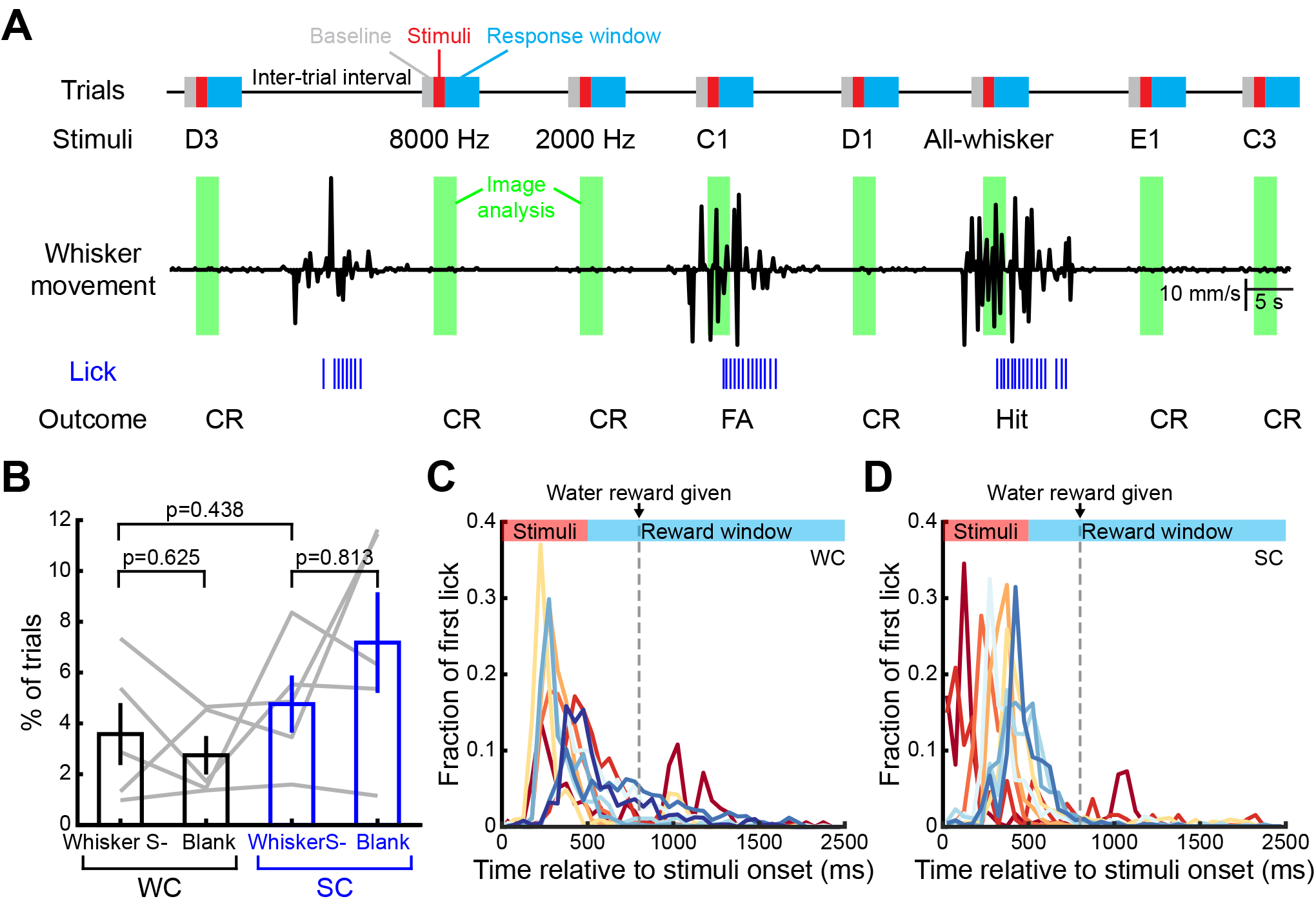
Mice withheld licking and whisking during whisker S− trials, and responded to cue but not delivery of water reward. Related to figure 1. **(A)** Contralateral whisker movement (black trace) and licking (blue raster lines) during an example behavioral segment containing 8 trials. Trials containing licks were not analyzed. CR: correct rejection; FA: false alarm. **(B)** Percentages of whisker S− trials and blank trials that contained whisker movement. Each gray line represents one mouse. Error bars: SEM. Statistics: Wilcoxon signed-rank test. **(C-D)** The timing of first lick in S+ trials for individual mice. Each line represented fractions of the beginning of lick bouts relative to stimuli onset in 50 ms time bins, averaged across all imaging sessions of one mouse. WC mice had 85.6% and SC mice had 93% of their first licks before reward delivery (p=0.497, rank-sum).

**Supplementary figure 2.**
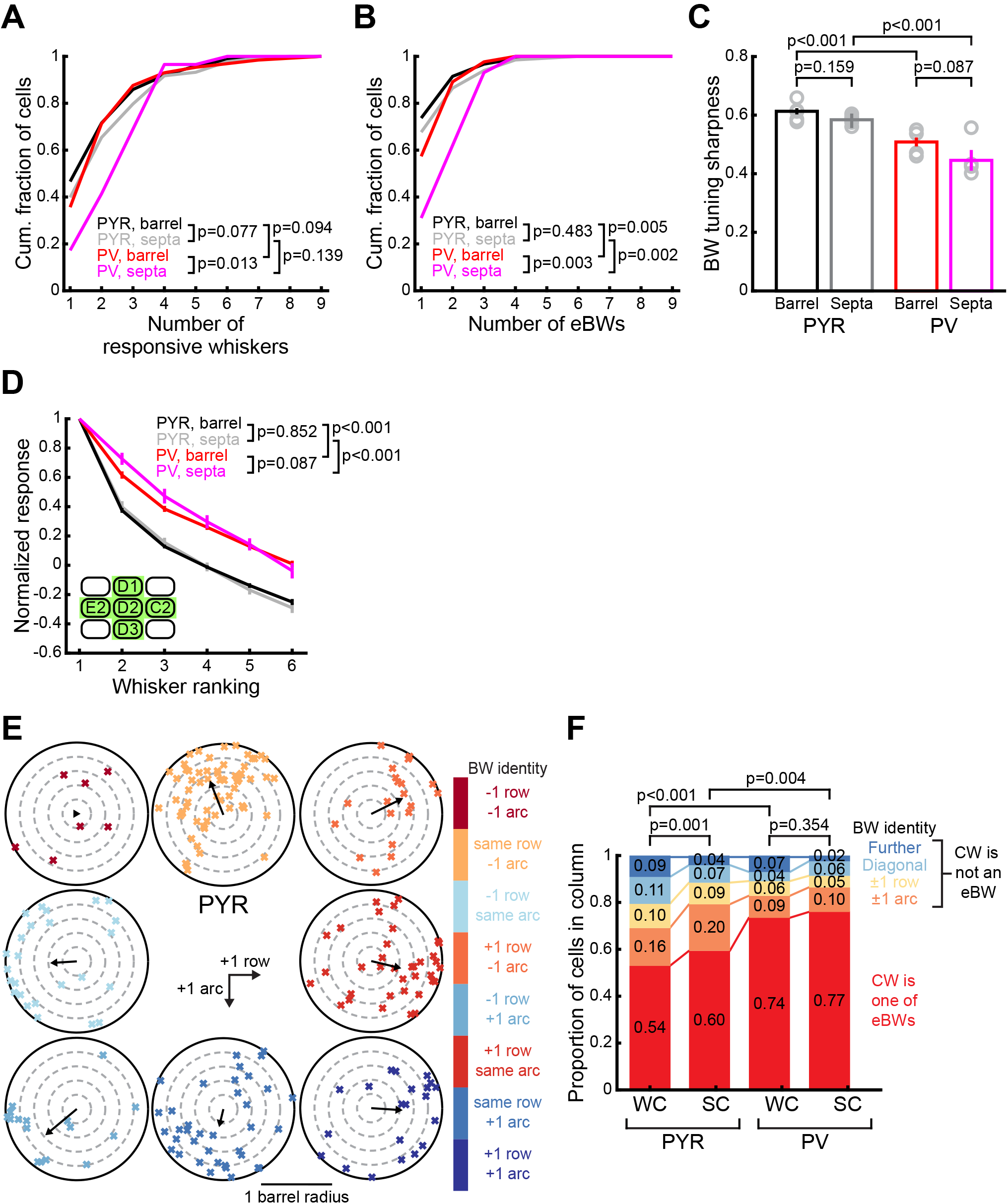
PYR and PV neuron tuning analyzed separately for barrel- and septa-related cells and whisker map topography. Related to figure 1 and 2. **(A-D)** L2/3 neurons were classified as being located above L4 barrels or L4 septa, and their tuning properties were analyzed separately. **(A)** Number of whiskers that evoked significant responses per cell. Statistics: χ^2^. **(B)** Number of eBWs for each neuron. Statistics: χ^2^. **(C)** Mean tuning sharpness of responsive neurons. Gray circles show the mean for each animal. Statistics: rank-sum. **(D)** Mean rank-ordered tuning curve, after ranking stimuli within cells. Only cells located in green columns (inset) were included. Statistics: unbalanced two-way ANOVA. These results show that barrel- and septa-related PYR cells have largely similar tuning precision, that PV tuning is broader than PYR tuning in both compartments, and that PV cells overlying septa have slightly broader tuning than PV cells overlying barrels, by some measures. In all panels, error bars are SEM. **(E)** Location of PYR neurons tuned to each of the 8 whiskers surrounding a reference whisker. Only cells located inside the reference whisker column were shown. The BWs of cells were color-coded, and the location of cells were presented as in Fig. 2D-E. The black circle indicated the border of reference whisker column. Arrows show circular mean of cell positions. **(F)** Tuning is also heterogeneous when defined by eBWs. Because of statistical uncertainty in identifying the BW from limited stimulus repetitions, we also characterized whisker map topography using eBWs, defined as all whiskers that evoked a mean ΔF/F that was statistically indistinguishable from the BW (see Methods). In (F), a neuron was considered tuned for the CW if the CW was among its eBWs. Panel showed fraction of responsive neurons whose eBWs included the CW (red), or that responded to another whisker significantly more than to the CW (other colors). Statistics: χ^2^.

**Supplementary figure 3.**
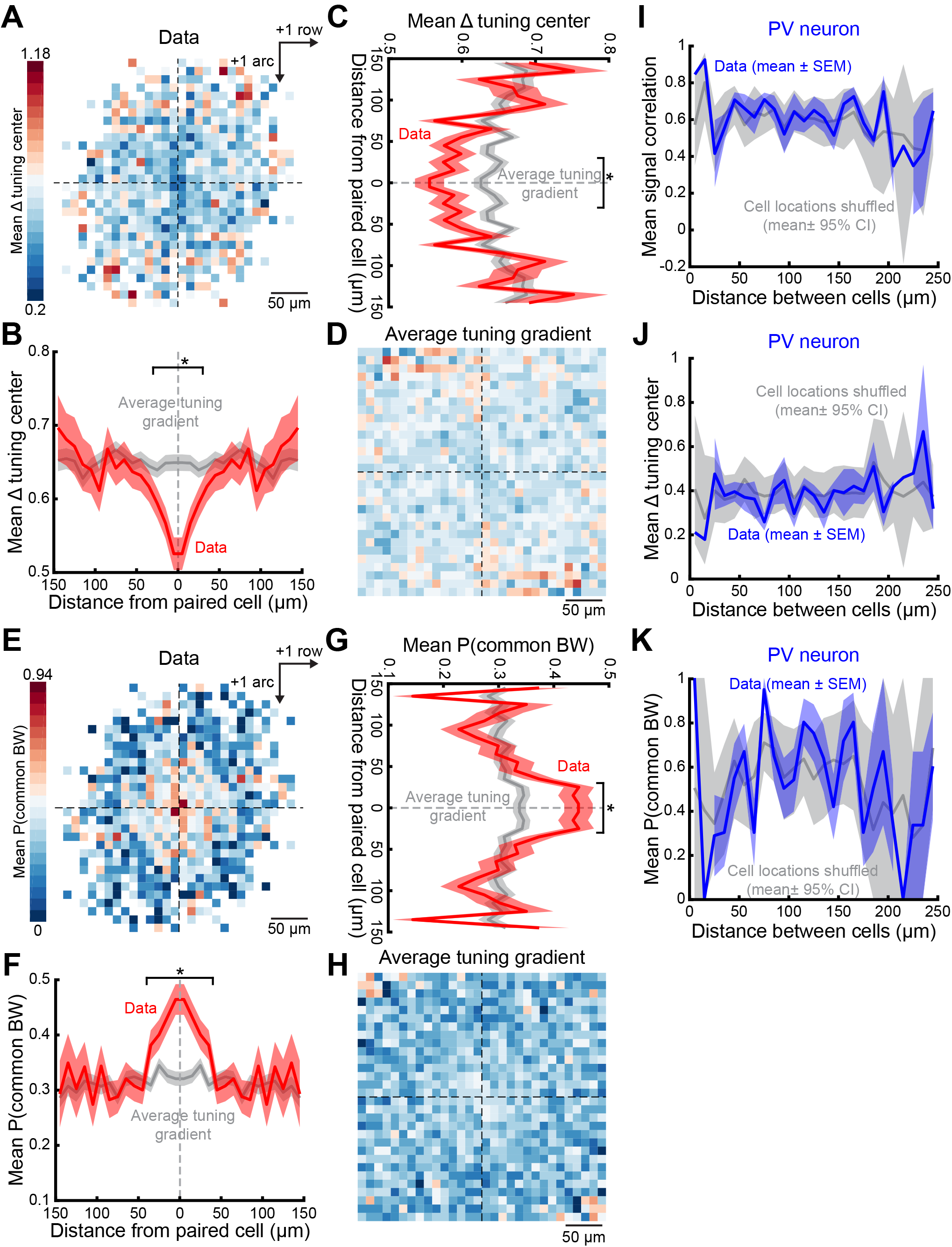
PYR neurons within each whisker column form local tuning clusters, but no local clustering for PV neurons. Related to figure 3. **(A-D)** Local tuning clusters defined by similarity in center-of-mass of whisker tuning. **(A)** Mean difference in tuning center-of-mass for pairs of co-columnar PYR neurons, as a function of 2-D spatial position of cells. For each neuronal pair, one cell is positioned at the center, and the tuning difference between the cells is assigned to the spatial bin at the location of the other cell. Color shows the mean tuning difference within each 10 µm spatial bin with cell numbers in each bin ≥ 5 cells. **(B-C)** Mean projection of data in (A) along row (B) or arc (C) dimension. *, significant difference within each bin relative to predicted column-average tuning gradient (gray). All shadings: SEM. **(D)** Predicted tuning gradient in the same coordinate system as (A), with local clustering removed by projecting cells across imaging fields and mice. See Methods. **(E-H)** Local tuning clusters defined by probability that two neurons share the same BW. (E) Probability that pairs of co-columnar PYR neurons have the same BW, as a function of 2-D spatial position of the cells. Plotted as in (A). Color shows the mean probability. **(F-G)** Mean projection of data in (E) along row (F) or arc (G) dimension. Plotted as in (B-C). *, significant difference within each bin relative to predicted column-average tuning gradient (gray). All shadings: SEM. **(H)** Predicted tuning gradient defined by the probability that two neurons share the same BW, after eliminating local clustering. Calculated and plotted as in (D). **(I-K)** Mean signal correlation (I), difference in tuning center-of-mass (J), and probability of sharing the same BW (K) for co-columnar PV pairs, as a function of distance between neuron pairs. Blue: observed data. Gray: spatially shuffled neurons. Plotted as in figure 3A,G,H, respectively. No local clustering was detected among PV neurons.

**Supplementary figure 4.**
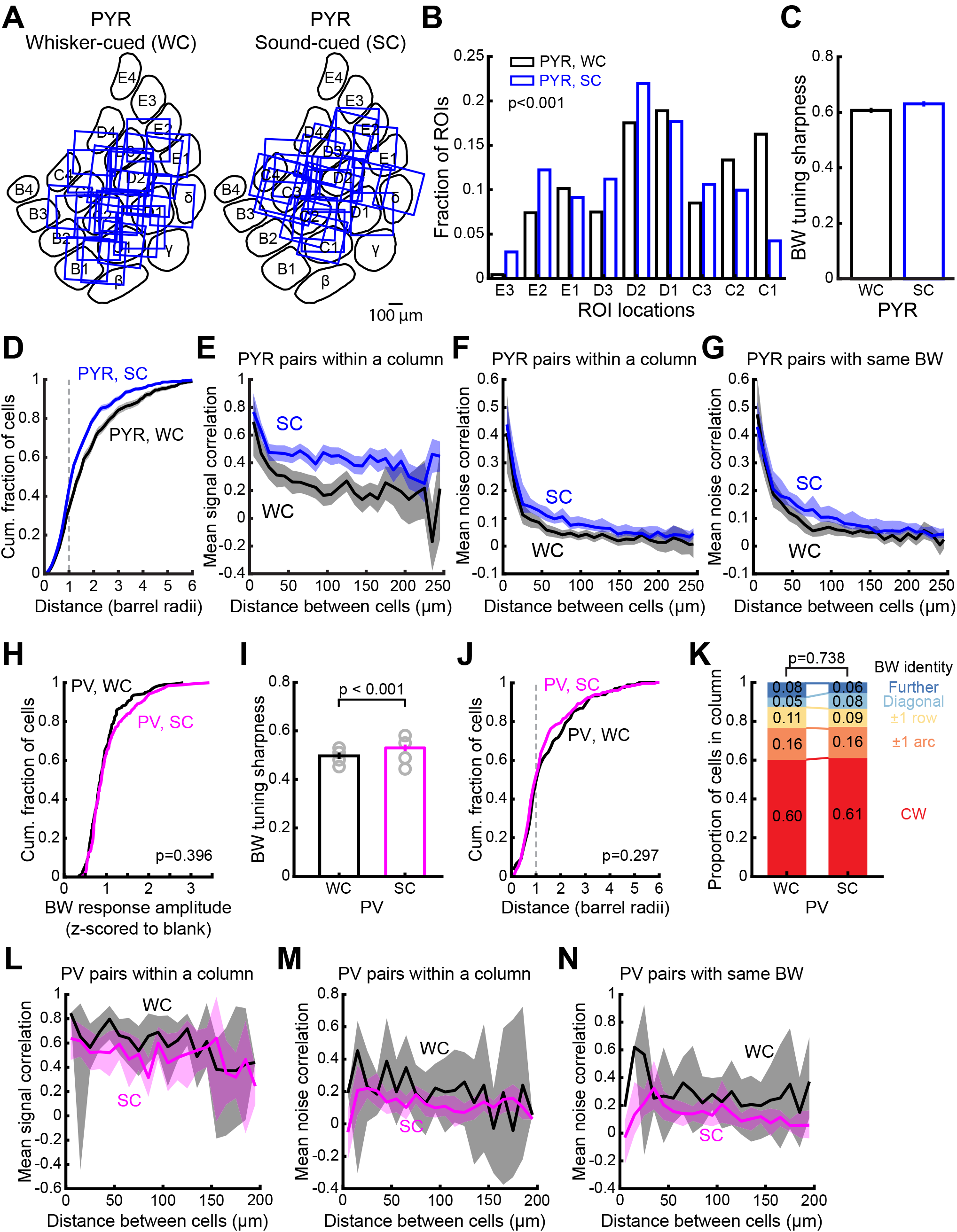
Whisker map differences between whisker-cued and sound-cued tasks. Related to figure 4. **(A)** PYR cell imaging field locations in WC and SC mice. **(B)** Anatomical columnar locations of all imaged PYR neurons, showing modest mismatch between WC and SC populations. Statistics: χ^2^. **(C-G)** Functional properties of PYR neurons in WC and SC mice compared after spatial sub-sampling to achieve exactly matched columnar distribution. See Methods. **(C)** Mean PYR tuning sharpness. **(D)** Cumulative distribution of distance from responsive PYR cells to their BW column center. **(E-F)** Mean signal (E) and noise (F) correlations for pairs of PYR neurons within the same column, averaged within each distance bin, as a function of distance between neuron pairs. **(G)** Same as (F), but for PYR pairs with the same BW. The mean correlations after resampling (panels E-G) were replotted in **Fig. 4I-K** as dashed lines. **(H-N)** PV neurons in WC and SC mice show similar tuning and topography. **(H)** Normalized BW response magnitude for WC and SC PV cells. Statistics: KS. **(I)** Mean tuning sharpness of responsive PV neurons. Gray circles show the mean for each animal. Statistics: rank-sum. **(J)** Cumulative distribution of distance from responsive PV cells to their BW column center. Statistics: KS. **(K)** Identity of BW for all responsive PV neurons within a column. Statistics: χ^2^. **(L-M)** Mean signal (L) correlation and noise (M) correlation for pairs of co-columnar PV neurons, averaged within each distance bin. **(N)** Same as (M), but for PV pairs with the same BW. In panel (L-M), raw data were sub-sampled to ensure exactly matched columnar distribution of PV neurons in WC and SC mice. In panels (I), error bars are SEM; error bars or shading in all other panels are 95% confidence intervals.

